# Antimicrobial activity of the quinoline derivative HT61, effective against non-dividing cells, in *Staphylococcus aureus* biofilms

**DOI:** 10.1101/2020.01.06.896498

**Authors:** C.J. Frapwell, P.J. Skipp, R.P. Howlin, E.M. Angus, Y. Hu, A.R.M. Coates, R.N. Allan, J.S. Webb

**Author notes:** denotes equal contribution (R.N. Allan, J.S. Webb). **Ethical Approval** Not required.

## Abstract

*Staphylococcus aureus* is an opportunistic pathogen responsible for a wide range of chronic infections. Disease chronicity is often associated with biofilm formation, a phenotype that confers enhanced tolerance towards antimicrobials, a trait which can be attributed to a dormant, non-dividing subpopulation within the biofilm. Development of antibiofilm agents that target these populations could therefore improve treatment success. HT61 is a quinoline derivative that has demonstrated efficacy towards non-dividing planktonic *Staphylococcus spp*. and therefore, in principal, could be effective against staphylococcal biofilms. In this study HT61 was tested on mature *S. aureus* biofilms, assessing both antimicrobial efficacy and characterising the cellular response to treatment. HT61 was found to be more effective than vancomycin in killing *S. aureus* biofilms (minimum bactericidal concentrations: HT61; 32 mg/L, vancomycin; 64 mg/L), and in reducing biofilm biomass. Scanning electron microscopy of HT61-treated biofilms also revealed disrupted cellular structure and biofilm architecture. HT61 treatment resulted in increased expression of proteins associated with the cell wall stress stimulon and *dcw* cluster, implying global changes in peptidoglycan and cell wall biosynthesis. Altered expression of metabolic and translational proteins following treatment also confirm a general adaptive response. These findings suggest that HT61 represents a new treatment for *S. aureus* biofilm-associated infections that are otherwise tolerant to conventional antibiotics targeting actively dividing cells.

## 1. Introduction

*Staphylococcus aureus* is commonly associated with chronic infections, particularly of the skin and soft tissue[1,2]. It will typically form sessile communities known as biofilms which are associated with increased tolerance to antimicrobials. Biofilms are highly heterogeneous, containing cellular sub-populations that are non-dividing and/or are metabolically inactive. As a large proportion of clinically administered antimicrobials target actively dividing cells this adopted quiescent state renders these antimicrobials ineffective, thus allowing biofilm bacteria to survive therapeutic intervention and contribute to chronic disease[3]. *S. aureus* has also evolved resistance towards common antimicrobials including β-lactams and glycopeptides, such as vancomycin (MRSA and VRSA, respectively)[4]; a trait which may be linked to increased horizontal gene transfer between bacteria residing in biofilms[5,6]. As a result of these tolerance and resistance mechanisms, ineffective treatment regimens can create environments that favour the selection of further antimicrobial resistance, making chronic infections even more difficult to treat[7]. As such, the development of novel antimicrobials targeting biofilm bacteria is highly desirable.

HT61 is a quinoline derivative that has demonstrated efficacy against both dividing and non-dividing planktonic cultures of *Staphylococcal spp*., with no detectable development of resistance[8–10]. HT61 preferentially binds to anionic staphylococcal membrane components, causing structural instability and cell depolarisation[8,10], and when used at a low concentration can also enhance the activity of neomycin, gentamicin and chlorhexidine against planktonic MSSA and MRSA[9]. Given its effectiveness against non-dividing *S. aureus*, HT61 represents an ideal candidate for targeting the dormant sub-populations present in *S. aureus* biofilms and could prove an effective treatment for biofilm-associated chronic infections.

In this study, we investigated the efficacy of HT61 against established *in vitro S. aureus* biofilms. We also utilised a quantitative label-free proteomic approach to identify changes in protein expression following treatment with sub-inhibitory and inhibitory concentrations in order to elucidate cellular processes linked to HT61’s mechanism of action.

## 2. Materials and Methods

### 2.1 Bacterial strains and growth conditions

*S. aureus* UAMS-1, a methicillin sensitive osteomyelitis isolate[11], was used for all experiments and grown in tryptic soy broth (TSB, Oxoid, UK) at 37 °C and 120 rpm (planktonic) or 50 rpm (biofilm). All inocula were performed with a starting density of 10^5^ cells ml^-1^. Biofilm cultures were performed in Nunc-coated 6 well plates, (Thermo-Fisher, UK) for susceptibility testing and proteomics, poly-L-lysine coated glass bottom dishes (MatTek, USA) for confocal laser scanning microscopy (CLSM), and glass cover slips in Nunc-coated 6 well plates for scanning electron microscopy (SEM). All biofilms were grown for 72 hours, with spent media replaced with fresh TSB every 24 hours.

### 2.2 Antimicrobial susceptibility testing

The efficacy of HT61 (Helperby Therapeutics) and vancomycin (Hospira Inc) was compared using planktonic and biofilm cultures of *S. aureus*. Cultures were treated with two-fold dilutions of each antimicrobial, (0.5 to 128 mg/L). Planktonic minimum inhibitory concentrations (MICs) were obtained using the broth microdilution method[9] and minimum bactericidal concentrations (MBCs) obtained by plating onto tryptic soy agar (TSA), prior to incubation at 37 °C and colony forming unit (cfu) enumeration. Biofilm MBCs were calculated as per Howlin *et al* (2015)[12]. Firstly, biofilms were grown as described in 2.1. Following 72 hours, spent media was replaced with medium containing either HT61 or vancomycin at the designated concentrations and biofilms were incubated for an additional 24 hours. The media was discarded and the biofilms rinsed twice with HBSS to remove non-adhered cells. Biofilms were detached using a cell scraper and suspended in 1 ml of HBSS. The suspensions were serially diluted, plated onto TSA and cfus were enumerated following a final 24 hour incubation. The MBCs were defined as the concentration that elicited a 99.9% reduction in viability.

### 2.3 Biofilm imaging

For CLSM, biofilms were grown in poly-L-lysine coated glass bottom dishes as described in 2.1. At 72 hours, media was replaced with TSB containing the biofilm MBC of HT61 or vancomycin (32 and 64 mg/L, respectively) and incubated for a further 24 hours. Prior to imaging, media was removed, biofilms rinsed twice with Hanks’ Balanced Salt Solution (HBSS; Sigma-Aldrich), then stained with 1 ml BacLight LIVE/DEAD (Life Technologies) (2 μl ml^−1^ of SYTO9 and Propidium Iodide) for 20 minutes. Biofilms were then washed once with HBSS then covered with 80% glycerol (v/v in HBSS) to prevent dehydration during imaging. Confocal images were acquired using an inverted Leica TCS SP8 CLSM and 63 X glycerol immersion lens with 1 μm vertical section slices. Images were analysed using the COMSTAT2.1 (Image J) image analysis package[13]. For SEM, biofilms were cultured on glass cover slips as described in 2.1. At 72 hours, media was replaced with TSB containing the biofilm MBC of HT61 (32 mg/L) and following a further 24 hour incubation, were then fixed in a solution of 3% glutaraldehyde, 0.1M sodium cacodylate (pH 7.2) and 0.15% alcian blue for 24 hours at 4 °C. A secondary fixative of 0.1 M sodium cacodylate (pH 7.2) and 0.1 M osmium tetroxide was performed for 1 hour. A final 1 hour fix was performed in 0.1 M sodium cacodylate (pH 7.2). Samples were dehydrated through an ethanol series (30:50:70:95:100:100%), critical point dried and sputter coating using a platinum-palladium alloy^14^. Imaging was performed using an FEI Quanta 250 SEM.

### 2.4 Protein Extraction

The cellular response of *S. aureus* to HT61 was determined using label-free ultra-performance liquid chromatography mass spectrometry^Elevated Energy^, (UPLC/MS_E_). Planktonic cultures were grown to stationary phase in TSB for 12 hours at 37 °C with either 0, 4 or 16 mg/L HT61 (sub-MIC and MIC, respectively) then centrifuged at 2500 *× g* for 15 minutes. Biofilms were cultured for 72 hours in Nunc-coated 6 well plates as described in 2.1. After 72 hours, media was replaced with TSB supplemented with 0, 4 or 16 mg/L HT61 and incubated for 12 hours at 37 ° C. Biofilms were rinsed twice using HBSS, scraped from the surface of the plate and resuspended into 1 mL HBSS. Cell suspensions were centrifuged at 2500 *× g* for 15 minutes, the supernatants removed, and the cell pellets rinsed twice with 0.1 M triethylammonium bicarbonate (TEAB). The cell pellets were then resuspended in 4 M Guanidine-HCl (in 0.1 M TEAB), then mechanically lysed using a TissueLyser LT (Qiagen) with Lysing Matrix B tubes (MP Biomedicals) at 50Hz for 20 x 30 second cycles with 30 second rest on ice between cycles). Proteins were precipitated in ice cold EtOH overnight at −20 °C, centrifuged at 12,000 *× g* for 10 minutes, then the pellets resuspended in 0.1 M TEAB and 0.1 % Rapigest SF Surfactant (Waters) prior to quantification using a Qubit fluorometer (Thermo-Fisher).

### 2.5 Sample Preparation for Mass Spectrometry

Dithiothreitol was added to 40 μg of each sample (final concentration 2.5 mM) and incubated at 56 °C for 1 hour. Iodoacetamide was added (final concentration 7.5 mM) and incubated at room temperature in the dark for 30 minutes prior to overnight digestion of samples at 37 ° C with 0.5 μg trypsin per sample. Trifluoroacetic acid (final concentration 0.5%) was added to each sample, before centrifugation (13, 000 *× g*, 10 minutes) and dried using a SpeedVac (Thermo-Fisher) to evaporate the solvent. Samples were resuspended in 0.5 % acetic acid, purified using a C18 solid phase extraction plate (Thermo-Fisher) according to manufacturer’s instructions and eluted into 80% acetonitrile and 0.5% acetic acid. Peptide extracts were lyophilised and stored at −20 °C until required. Peptide extracts were re-suspended in buffer A, (3 % acetonitrile, 0.1 % formic acid (v/v)) at a concentration of 0.25 μg/μL containing the internal digest standard, yeast enolase (Waters) at a final concentration of 0.25 fmol/μL.

Prepared samples were analysed using a Waters Synapt G2Si high definition mass spectrometer coupled to a nanoAcquity UPLC system. 4 μl of peptide extract was injected onto a C18 BEH trapping column (Waters) and washed with buffer A for 5 min at 5 μl/min. Peptides were separated using a 25 cm T3 HSS C18 analytical column (Waters) with a linear gradient of 3-50 % acetonitrile + 0.1 % formic acid over 50 minutes at a flow rate of 0.3 μl/min. Eluted samples were sprayed directly into the mass spectrometer operating in MS^E^ mode. Data were acquired from 50 to 2000 m/z with the quadrupole in RF mode using alternate low and elevated collision energy (CE) scans, resolution of 35,000. Low CE was 5 V and elevated CE ramp from 15 to 40 V. Ion mobility separation was implemented prior to fragmentation using a wave velocity of 650 m/s and wave height of 40 V. The lock mass Glu-fibrinopeptide, (M+2H)+2, m/z = 785.8426) was infused at a concentration of 100 fmol/μl at a flow rate of 250 nl/min and acquired every 60 sec.

### 2.6 Data processing

Raw data were processed using a custom package (Regression tester) based upon executable files from ProteinLynx Global Server 3.0 (Waters). The optimal setting for peak detection across the dataset was determined using Threshold inspector (Waters) and these thresholds were applied: low energy = 100 counts; high energy = 30 and a total energy count threshold of 750. Database searches were performed using regression tester and searched against the Uniprot *S. aureus* MN8 reference database (accessed 25/01/2018) with added sequence information for the internal standard Enolase. A maximum of two missed cleavages was allowed for tryptic digestion and the variable modifications were set to oxidation of methionine and carboxyamidomethylation of cysteine.

### 2.7 Data Analysis

All experiments were performed with a minimum of 3 biological replicates. Statistical analyses were performed using GraphPad Prism version 7.0d for Mac, Microsoft Excel and R version 3.6.0, ggplot2, and cowplot[14–16]. Significance was designated at *p* ≤ 0.05.

For mass spectrometry data, each data set was normalised to the top 200 most abundant proteins (per ng). Inclusion criteria for quantitative analysis and comparison were as follows; the protein must be present in all 3 biological replicates with a false discovery rate (FDR) ≤ 1% and sequence coverage ≥ 5%. Differential expression was categorised by an expression rate of ≥ 1.5 and ≤ 0.667 with *p* ≤ 0.05 using a one-tailed student t-test. Proteins were analysed using a combination of uniprot database searches (www.uniprot.org, accessed between 01/05/18 and 07/07/18) and gene ontology analysis using GeoPANTHER[17].

## 3. Results

### 3.1 HT61 is more effective than vancomycin towards *S. aureus* biofilms

The efficacy of HT61 and vancomycin towards planktonic and biofilm cultures of *S. aureus* was compared (Table 1). The planktonic MIC and MBC values for HT61 were 16 mg/L and 32 mg/L respectively in comparison to 4 mg/L for vancomycin. However, based on the definition of MBCs, HT61 was twice as effective as vancomycin at treating *S. aureus* biofilms with an MBC of 32 mg/L compared to 64 mg/L respectively. When viable counts were considered, HT61 consistently presented with improved killing of *S. aureus* UAMS-1 biofilms compared to vancomycin, over a range of concentrations. At the maximum concentration tested (128 mg/L), HT61 caused a further 1.3 log reduction in CFUs compared to vancomycin utilised at the same concentration (Figure 1).

**Figure 1:**
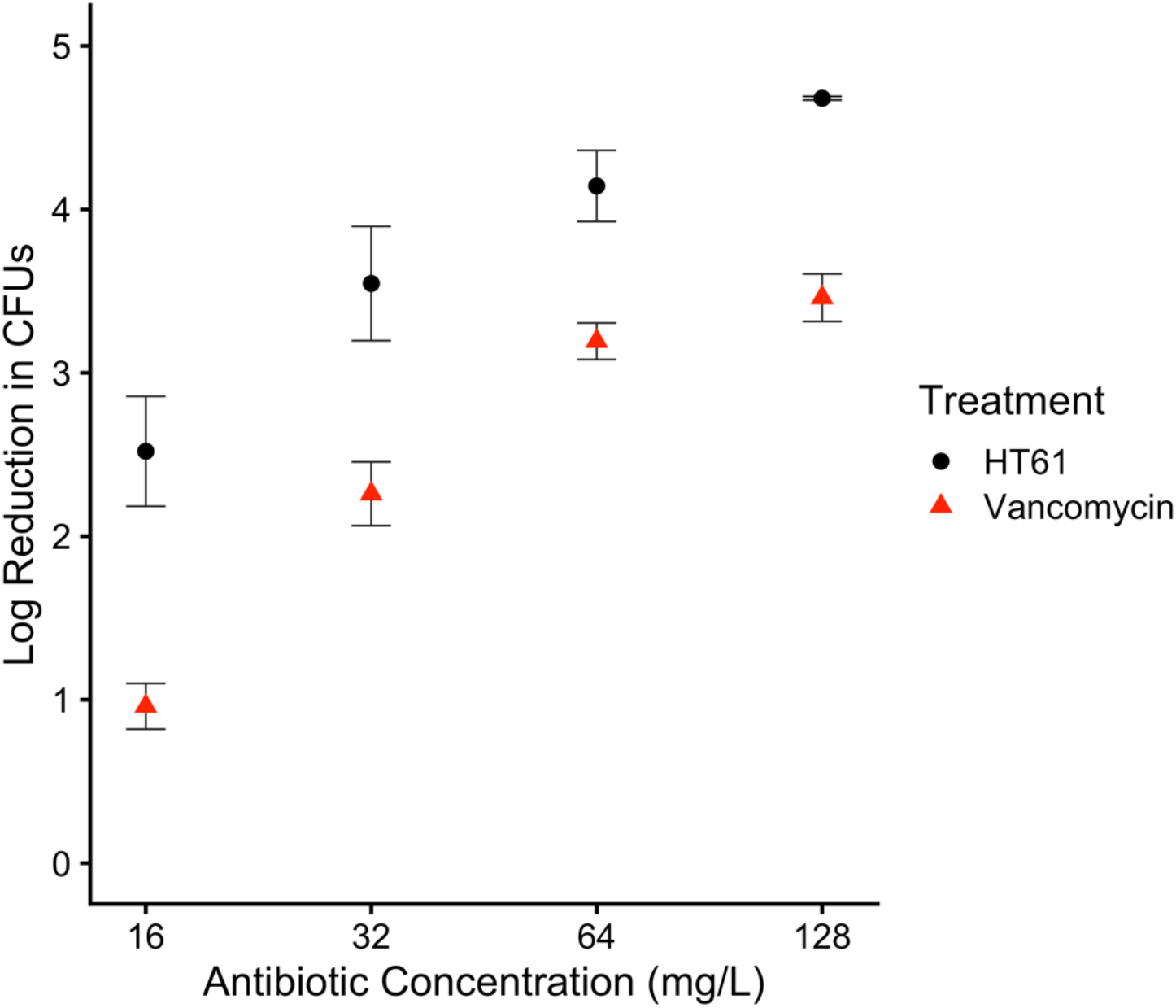
Log Reduction in *S. aureus* UAMS-1 viable counts of an established 72 hour biofilm following treatment with HT61 and vancomycin. HT61 consistently elicited a greater log reduction in CFU counts than vancomycin, demonstrating its potential as an antibiofilm agent. A higher value indicates a greater log reduction in CFUs. n = 3. Error bars indicate standard deviation.

**Table 1:** HT61 and vancomycin susceptibility of planktonic and biofilm cultures of *S. aureus* UAMS-1. Vancomycin was more effective than HT61 at treating planktonic cultures, but less effective at treating biofilm cultures (n = 3).

|  | MIC | MBC | Biofilm MBC |
| --- | --- | --- | --- |
|  | mg/L |  |  |
| HT61 | 16 | 32 | 32 |
| Vancomycin | 4 | 4 | 64 |

Notably, unlike vancomycin, which was less effective against biofilms (64 mg/L) compared to planktonic cultures (4 mg/L), HT61 was equally as effective towards planktonic and biofilm cultures (both 32 mg/L).

When treating 72 hour established *S. aureus* biofilms with their respective biofilm MBCs (32 mg/L HT61 and 64 mg/L vancomycin) for 24 hours they both had a similar effect on biofilm architecture, causing significant decreases in biofilm thickness and the surface area to volume ratio of live biomass (Figure 2A-C). However, a non-significant reduction in biofilm thickness and live biomass was observed with HT61 compared to vancomycin (Figure 2D and E, *p* > 0.05).

**Figure 2:**
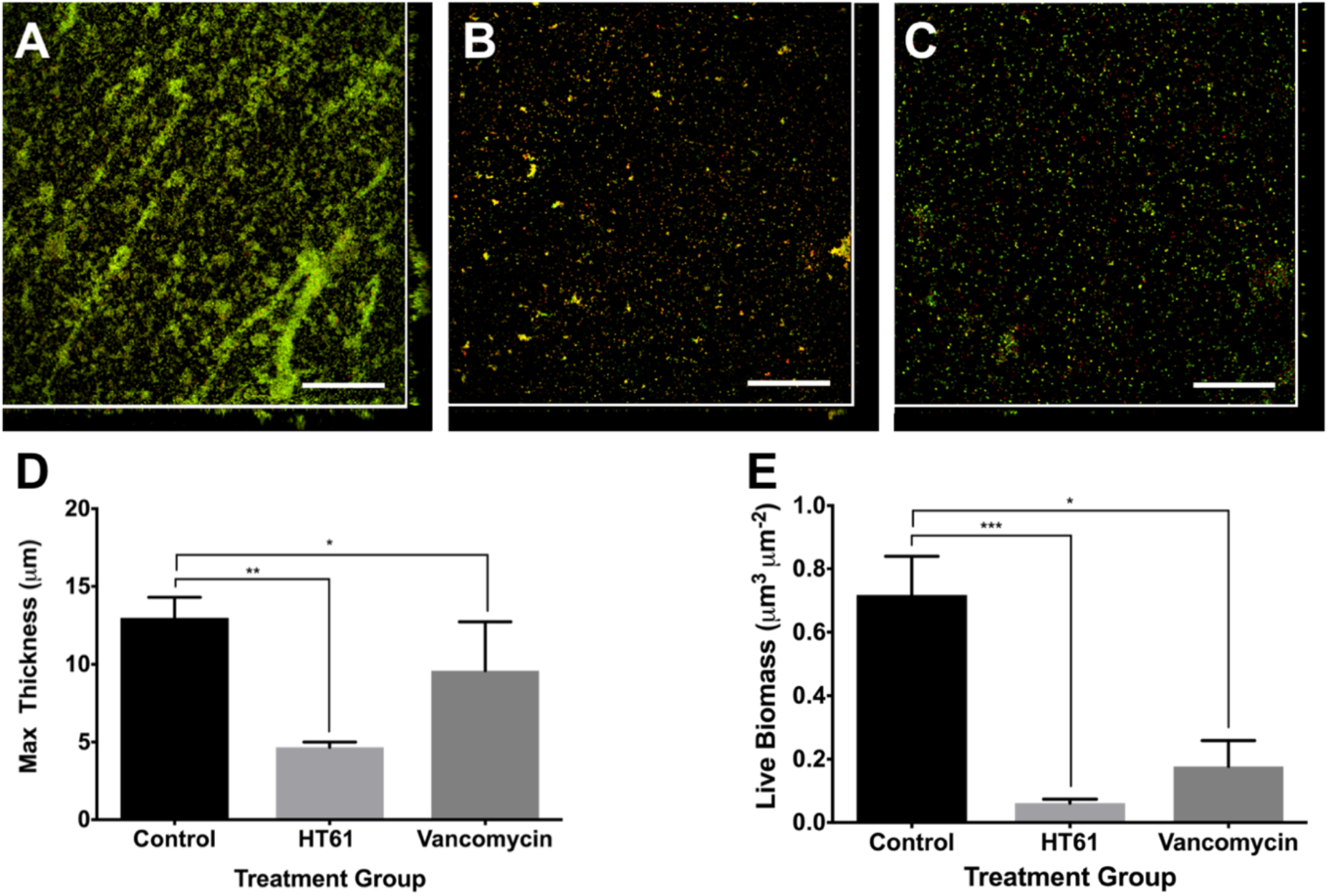
Representative confocal images of 96 hour *S. aureus* UAMS-1 biofilms that were **(A)** untreated, **(B)** treated with 32 μg ml^−1^ HT61 for 24 hours, and **(C)** treated with 64 μg ml^−1^ vancomycin for 24 hours. Biofilms were stained with 1 ml of BacLight Live/Dead (2 μl ml^−1^ SYTO9 and Propidium Iodide) for 20 minutes. Live biomass is depicted in green, while dead biomass is red. Scale bars = 50 μm. COMSTAT analysis was performed to assess the effect of treatment on **(D)** maximum biofilm thickness and **(E)** live biofilm biomass. Error bars represent standard error of the mean (n = 12). Statistical significance determined by Kruskal-Wallis Test and post-hoc Dunn’s comparisons * (P ≤ 0.05), ** (P ≤ 0.01), *** (P ≤ 0.001).

### 3.2 HT61 treatment of *S. aureus* biofilms disrupts the cell envelope

SEM was used to examine the effect of 24 hour HT61 treatment on individual *S. aureus* cells within a 96 hour biofilm. Prior to treatment biofilms were comprised of dense cell aggregates associated with an extracellular polymeric substance (EPS). Cellular morphology was typical with no obvious signs of stress or damage (Figure 3 A-C). Following treatment with 32 mg/L HT61 both the size and quantity of biofilm aggregates were reduced. Severe damage to individual cellular structures was also observed, indicative of resultant stress and lysis (Figure 3D-F).

**Figure 3:**
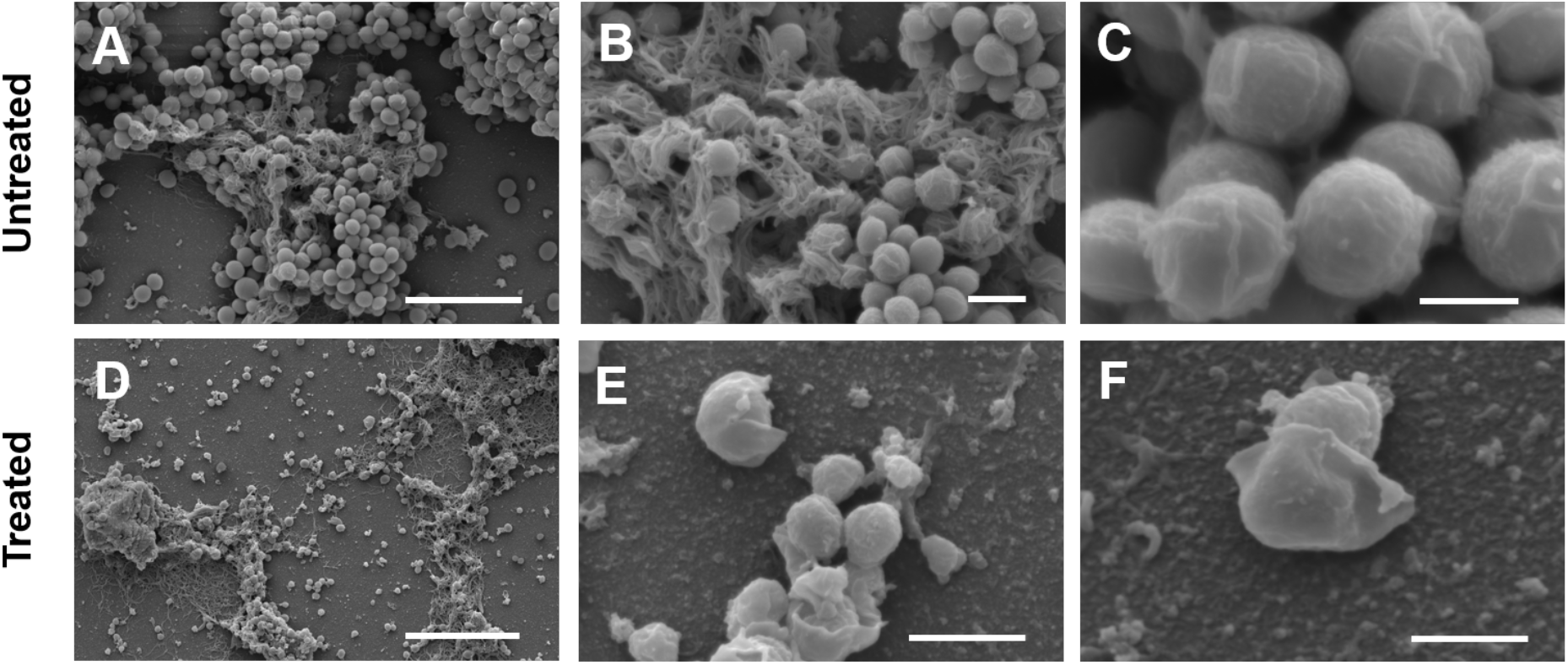
Representative SEM micrographs of 96 hour *S. aureus* UAMS-1 biofilms that were either untreated **(A-C)** or treated with the 32 mg/L HT61 for 24 hours **(D-F)**. Scale bars: A and D = 5 μm, B and E = 1 μm, C and F = 0.5 μm.

### 3.3 *S. aureus* biofilm growth is associated with large-scale metabolic changes including increased glucose and arginine catabolism

Mass spectrometry of planktonic and biofilm cultures of *S. aureus* identified a total of 1,448 proteins. 462 proteins met the inclusion criteria for quantitative analysis, with 60 (13.0%) increased in expression and 89 (19.3%) decreased in expression (Figure 4, complete list available in Table S1). Of the proteins that were differentially expressed the majority were involved in metabolic processes, representing 46.7% of proteins increased in expression and 25.8% of proteins decreased in expression (see Table S2). All proteins identified as part of the TCA cycle were significantly upregulated, (9/9, average 4.99-fold increase, with CitZ citrate synthase upregulated 11.95-fold). Conversely, all identified proteins associated with fatty acid metabolism (5/5) were downregulated. Proteins associated with glycolysis/gluconeogenesis were also affected during biofilm growth with 26% (5/19) upregulated, 22% (4/19) downregulated and 58% (11/19) commensurate to planktonic expression. Due to the upregulation of the TCA cycle, this expression profile likely corresponds to an increase in glycolytic activity, rather than gluconeogenesis. Proteins important for amino acid metabolism were also differentially expressed in biofilms, with 50% (6/12) upregulated and 25% (3/12) downregulated. Associated with this was the upregulation of components associated with one carbon/folate metabolism (which is linked to amino acid and nucleotide metabolism), with 80% (4/5) upregulated and 20% (1/5) downregulated. These data suggest that amino acid requirements are significantly altered during biofilm growth. Increased expression of ArcA, ArcB and ArcC infers upregulation of the arginine deiminase (ADI) pathway, which revolves around the catabolism of L-arginine, resulting in 1 mol ATP and is important for growth under anaerobic conditions[18]. The remaining metabolic proteins encompassed a number of functions but no particular processes were enriched. 79% (51/65) of these proteins were not differentially expressed suggesting that they are important in both the planktonic and biofilm phenotypes.

**Figure 4:**
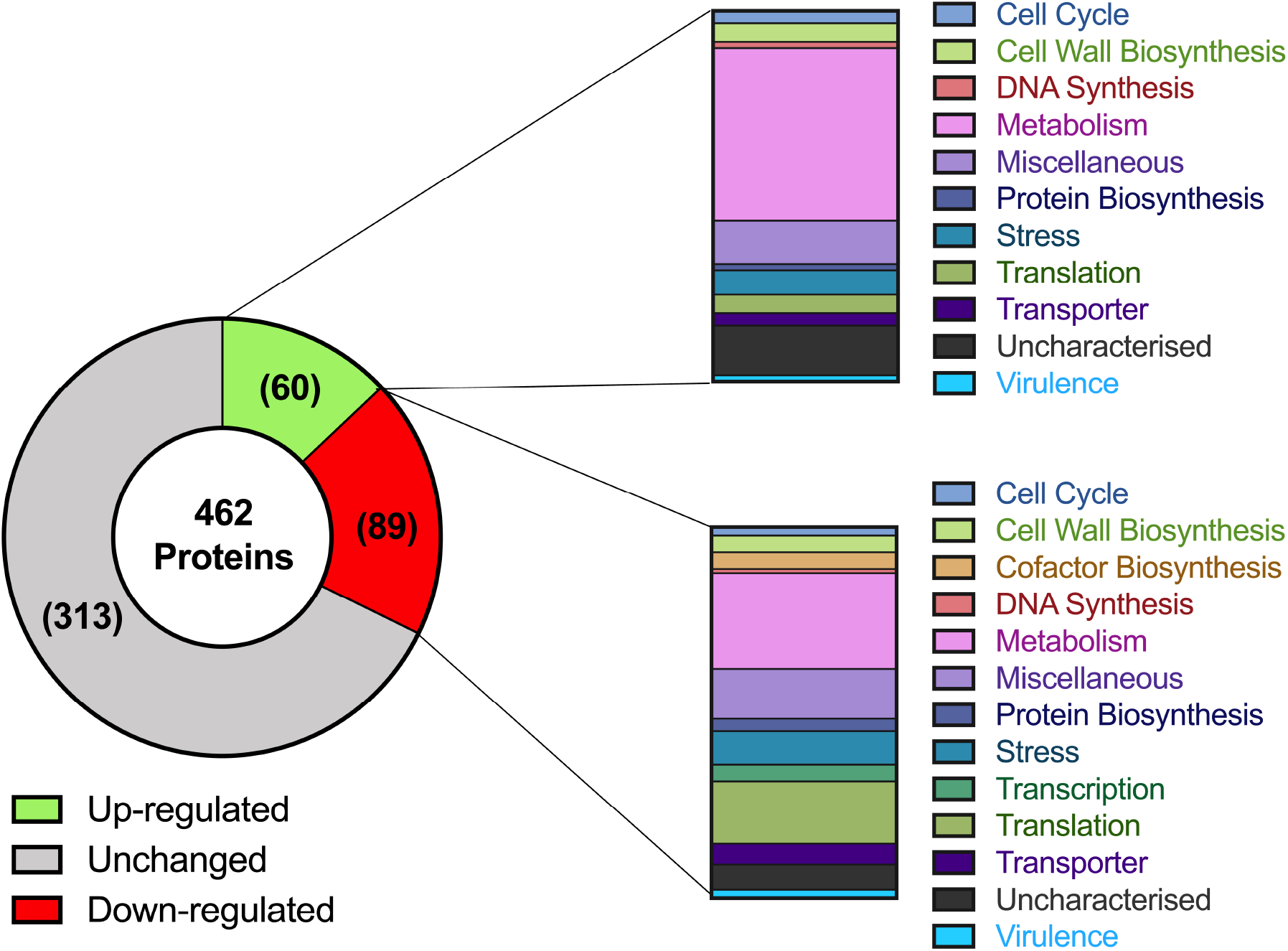
Overview of differential protein expression between planktonic and biofilm cultures of *S. aureus*. Number of proteins identified for each category is displayed in brackets. For quantitative analysis the following selection criteria were set: proteins present in all three biological replicates, FDR of ≤ 1%, and sequence coverage of ≤ 5%. Proteins were classed as differentially expressed at ≥ 1.5 for increased expression or ≤ 0.667 for decreased expression, p ≤ 0.05. Functional categories were assigned using GeoPANTHER and UniProt database searches.

### 3.4 HT61 Treatment results in increased expression of the Cell Wall Stress (CWS) stimulon and *dcw* cluster

The proteomic response of planktonic and biofilm cultures of *S. aureus* was compared following treatment with HT61 at either sub-MIC (4 mg/L) or MIC (16 mg/L) concentrations (Table 2). HT61 treatment resulted in the differential expression of proteins involved in a variety of functions including cell wall biosynthesis, DNA synthesis, and metabolism (see Tables S3 and S4).

**Table 2:**
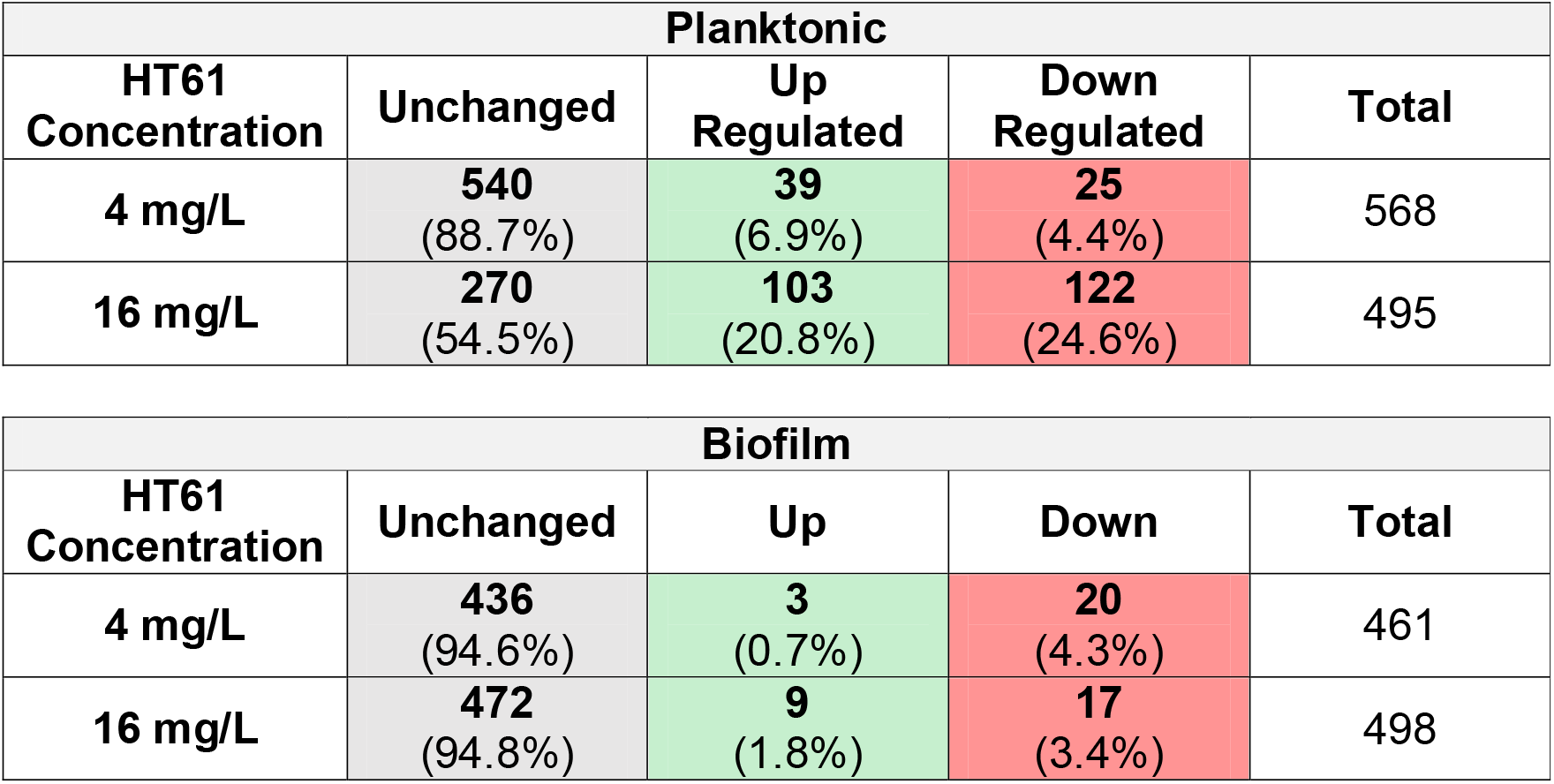
Summary of differential protein expression between untreated, sub-MIC (4 mg/L), and MIC (16 mg/L) treated *S. aureus* planktonic and biofilm cultures. Inclusion criteria for quantitative analysis and comparison was set at 3 peptide matches, false discovery rate (FDR) ≤ 1%, sequence coverage ≥ 5%, with p ≤ 0.05.

For planktonic cultures treated with sub-MIC HT61 (4 mg/L), two cell wall biosynthesis associated proteins required for the incorporation of D-glutamate into cell wall peptidoglycan[19], MurD and MurI, were upregulated (Table 3). Increasing the concentration of HT61 from 4 mg/L to 16 mg/L led to upregulation of 93% (14/15) of proteins associated with cell wall biosynthesis, including 6 components of the mur ligase pathway (MurACDEFI, 2.63 mean fold increase), FemA-like protein and FemB, which are required for peptidoglycan crosslinking (2.53 mean fold increase) and a 2.19 fold upregulation of VraR, the regulator of the CWS stimulon[20].

**Table 3:** Differentially expressed proteins associated with the *dcw* and cell wall stimulon in *S. aureus* following treatment of planktonic cultures with HT61. Expression ratios reflect changes in expression between untreated cultures and those treated with either sub-MIC (4 mg/L) or MIC (16 mg/L) concentrations of HT61. Differential expression in biofilms indicated in brackets. Differential expression is defined as a fold change ≥ 1.5 for upregulation (green cells) and ≤ 0.667 for down regulation (red cells). Grey cells indicate no change in expression. Empty cells – proteins not identified.

|  | Accession Number | Protein Name | Gene | Expression Ratio |  |
| --- | --- | --- | --- | --- | --- |
|  |  |  |  | Sub-MIC | MIC |
| Cell Cycle | A0A0E1X830_STAAU | Cell division protein FtsA | *ftsA* | 1.38 | 1.66 |
|  | A0A0E1X718_STAAU | GTPase Obg | *cgtA* | 1.30 | 3.04 |
|  | A0A0E1X5J2_STAAU | Cell cycle protein GpsB | *gpsB* | 1.10 | 0.20 |
|  | A0A0E1XAY0_STAAU | 60 kDa chaperonin | *groL* | 1.13 | 0.29 |
|  | A0A0E1XGT1_STAAU | DivIVA domain protein | HMPREF0769_12587 | 1.05 | 0.29 |
|  | A0A0E1X4P6_STAAU | Trigger factor | *tig* | 1.01 | 0.34 |
| Cell Wall Biosynthesis | A0A0E1XHI9_STAAU | DltD central region | *dltD* | 1.78 | 2.51 |
|  | A0A0E1X5R6_STAAU | FemAB family protein (FemA) | HMPREF0769_12373 (*femA*) | 1.05 | 1.82 |
|  | A0A0E1XIT0_STAAU | UDP-N-acetylglucosamine 1-carboxyvinyltransferase | *murA1* | 0.98 | 2.05 |
|  | A0A0E1XAN0_STAAU | UDP-N-acetylglucosamine 1-carboxyvinyltransferase | *murA2* | 1.12 | 2.83 |
|  | A0A0E1X4D8_STAAU | UDP-N-acetylmuramate--L-alanine ligase | *murC* |  | 2.40 |
|  | A0A0E1X8P8_STAAU | UDP-N-acetylmuramoylalanine--D-glutamate ligase | *murD* | 1.84 | 3.43 (1.59 Biofilm) |
|  | A0A0E1X6V3_STAAU | UDP-N-acetylmuramoyl-L-alanyl-D-glutamate--L-lysine ligase | *murE* | 1.05 | 1.76 |
|  | A0A0E1XIV1_STAAU | UDP-N-acetylmuramoyl-tripeptide--D-alanyl-D-alanine ligase | *murF* | 1.33 | 2.31 |
|  | A0A0E1X8U4_STAAU | Glutamate racemase | *murI* | 1.52 | 3.62 |
|  | A0A0E1XKB3_STAAU | Ribulose-5-phosphate reductase | *tarJ* | 1.12 | 2.58 |
|  | A0A0E1XJG3_STAAU | Response regulator protein VraR | *vraR* |  | 2.19 |
|  | A0A0E1X974_STAAU | Mur ligase middle domain protein | HMPREF0769_11280 | 1.32 | 2.67 |
|  | A0A0E1X785_STAAU | D-alanine--D-alanyl carrier protein ligase | *dltA* | 1.15 | 1.92 |
|  | A0A0E1XG48_STAAU | Aminoacyltransferase FemB | *femB* | 0.99 | 3.24 |
|  | A0A0E1X6S7_STAAU | Mannosyl-glycoprotein endo-beta-N-acetylglucosaminidase | HMPREF0769_12730 | 0.89 | 0.63 |
| DNA<br>Maintenance/Synthesis | A0A0E1XAS7_STAAU | ATP-dependent DNA helicase | *pcrA* | 1.31 | 3.07 (2.13 Biofilm) |
|  | A0A0E1X928_STAAU | DNA ligase | *ligA* | 1.29 | 1.73 |
|  | A0A0E1XAK8_STAAU | Chromosomal replication initiator protein DnaA | *dnaA* | 2.07 | 2.90 |
|  | A0A0E1XB29_STAAU | DNA polymerase III subunit gamma/tau | *dnaX* | 1.60 | 2.07 |
|  | A0A0E1X6I5_STAAU | DNA polymerase I | *polA* | 1.37 | 1.51 |
|  | A0A0E1XAK2_STAAU | DNA gyrase subunit A | *gyrA* | 1.12 | 1.55 |
|  | A0A0E1X7H6_STAAU | DNA topoisomerase 4 subunit B | *parE* | 1.30 | 3.34 |
|  | A0A0E1XFV3_STAAU | DNA-binding protein HU | *hup* | 0.91 | 0.33 |
|  | A0A0E1X9G8_STAAU | Nucleoid-associated protein HMPREF0769_10004 | HMPREF0769_10004 |  | 0.15 |

DNA synthesis in planktonic cultures was also affected by HT61 treatment (Table 3). Sub-inhibitory treatment increased expression of DnaA and DnaX, indicating a general increase in DNA synthesis (mean 1.84 fold increase). Increasing the concentration of HT61 to 16 mg/L led to the increased expression of more proteins associated with DNA maintenance. Notably, three of the upregulated proteins (PcrA, GyrA and ParE) possess DNA helicase activity, responsible for the relaxation of DNA super-coiling. Alongside the increased expression of DNA synthesis/maintenance components there was differential expression of cell cycle associated proteins, with two upregulated (FtsA and Obg, mean 2.35 fold increase) and four downregulated (GpsB, GroL, Tig and DivlVA domain protein, mean 0.28 fold decrease). As well as being part of the CWS, a number of the differentially expressed cell wall biosynthesis components, DNA synthesis genes and cell cycle components comprise a segment of the division cell wall, *dcw* cluster, a family of genes that are vital for maintaining cell shape and integrity[21,22].

Biofilms treated with HT61 presented with a similar, albeit more muted response (Table 2). When treated with HT61 at 16 mg/L increased expression was observed for both MurD (1.59 fold) and PcrA (2.13 fold), similar to planktonic cultures (Table 3). Unlike HT61 treated planktonic cultures, proteins associated with the cell cycle were not differentially expressed in HT61 treated biofilms.

### 3.5 HT61 treatment affects cellular metabolism and translation

Sub-MIC HT61 treatment of planktonic cultures was associated with increased expression of components of the ADI pathway (ArcA, ArcC) and ArgF (1.93 mean fold increase). However, increasing the HT61 concentration resulted in these proteins returning to basal expression, but led to other significant changes in cellular metabolism. These included decreased expression of two TCA cycle components (succinate dehydrogenase SdhA, and citrate synthase CitZ) and six proteins associated with glycolysis/gluconeogenesis. Conversely, there was upregulation of proteins linked to fatty acid metabolism (80%, 4/5), as well as other miscellaneous metabolic processes (see Table S3). Curiously, when utilised at its MIC, HT61 treatment of planktonic cultures led to the upregulation of 5 proteins linked to tRNA modification. All proteins were upregulated more than two-fold, with MnmG and YqeV upregulated 3.65 and 7.32 fold, respectively. tRNA modifications alter tRNA base specificity and alter translational output[23].

## 4. Discussion

The dormant populations of bacteria residing within biofilms are a major contributing factor towards the enhanced antimicrobial tolerance associated with this phenotype, and also serve as a potential driver for the development of AMR^3^. Targeting these populations therefore offers a route to treating biofilm-associated chronic infections and reducing the spread AMR. Previous studies have shown that the quinoline derivative, HT61, was effective in treating non-dividing planktonic staphylococcal *spp*., thereby making it an ideal candidate for the treatment of *S. aureus* biofilms^8-10^.

To determine whether HT61 would be more effective than an antibiotic that targets actively dividing cells we compared its activity to vancomycin, for which the mechanism of action necessitates active cell wall turnover[24]. Whilst vancomycin was more effective than HT61 against planktonic *S. aureus* its efficacy was reduced against the biofilm phenotype, evidencing the enhanced tolerance which can be attributed to the presence of a dormant subpopulation. HT61, however, was more effective than vancomycin at treating established *S. aureus* biofilms, confirming that its activity against non-dividing cells confers an advantage against this phenotype. Confocal microscopy demonstrated that HT61 reduced both biofilm biomass and viability, with SEM imaging also revealing that HT61 treatment disrupts the outer layers of the cell, (cell wall and cell membrane), as previously documented[8,10].

Label-free UPLC/MS^E^ was used to quantify the cellular response of planktonic and biofilm cultures of *S. aureus* to HT61. Comparing baseline protein expression profiles of both phenotypes in the absence of treatment indicated that metabolic changes were most abundant. Notably, the transition to the biofilm phenotype was associated with increased expression of the TCA, glycolytic and ADI pathways. Activation of the TCA cycle and glycolysis (particularly decreased expression of lactate dehydrogenase, see Table S2), implies increased aerobic metabolism. Conversely, activation of the ADI pathway implies increased anaerobic and anoxic metabolism as the cells attempt to generate energy via the conversion of L-arginine to L-citrulline[18]. While these processes appear conflicting, their simultaneous activation highlights the differing energy requirements of biofilm cells as well as the stark physiological heterogeneity found in a *S. aureus* biofilm after only 72 hours of growth. Co-current activation of these metabolic processes also corroborates previous proteomic and transcriptomic studies of *S. aureus* biofilms[25,26].

The response of *S. aureus* to HT61 involved the differential expression of proteins linked to several cellular functions and provided insight into HT61’s mechanism of action. Notably, HT61 treatment was associated with upregulation of proteins linked to peptidoglycan and cell wall biosynthesis, as well as the cell cycle and DNA synthesis. These proteins form part of the CWS stimulon, activated following stress to the cell envelope, and the *dcw* cluster, responsible for maintaining cell shape and integrity[22,27]. HT61 has been shown to preferentially bind to anionic phospholipids in the *S. aureus* cell membrane, in a manner similar to the lipopeptide antimicrobial, daptomycin[10,28,29]. Daptomycin inserts into the cell membrane, leading to alterations in membrane curvature, potassium efflux and membrane depolarisation[28,29]. Daptomycin mediated membrane curvature has been shown to directly impair cell wall synthesis by impairing the function of the cell wall biosynthesis protein, MurG[30]. Transcriptional profiling has also shown that daptomycin can also upregulate components of the cell wall stimulon, suggesting a secondary mechanism of action and/or interactions with the associated components[31]. Altered expression of the *dcw* cluster has also been documented in biofilms of *Haemophilus influenzae* following D-methionine treatment, contributing to altered cell morphology[21]. It is possible that HT61 functions in a similar manner to these examples, either by directly interfering with cell wall biosynthesis machinery or by placing undue levels of stress directly on the cell membrane, thereby interfering with the cell wall machinery. A related observation was that HT61 treatment led to the upregulation of several DNA helicases in planktonic cultures. While this could be a result of general DNA stress, it is possible that HT61 is moonlighting as a DNA gyrase inhibitor, similar to other quinoline-like antimicrobials, such as ciprofloxacin[32]. Finally, metabolic processes were generally decreased following HT61 treatment. This may be an attempt by the cell to limit HT61 damage, similar to the proteomic response of MSSA to oxacillin[33]. The proteomic response of established biofilms to treatment with the planktonic MIC concentration of HT61 was reduced compared to the response of planktonic cultures. This suggests that the remaining biofilm population, by nature of their dormant state and distinct expression profiles, are not responding to treatment and are likely to be susceptible to the confirmed MBC concentration of HT61 (32 mg/L).

In conclusion, we have demonstrated that HT61 is more effective than vancomycin in treating *S. aureus* biofilms by virtue of its ability to target dormant subpopulations, and therefore represents a potential treatment for chronic *S. aureus* biofilm infections. Using a quantitative proteomic approach, we have also shown that HT61 can influence the expression of the CWS stimulon and *dcw* cluster. By identifying cellular processes that altered following HT61 treatment, we suggest that they may provide novel antimicrobial targets and could inform the development of future antimicrobial compounds and therapeutic strategies.

## Supporting information

Supplementary Proteomics Data

## Acknowledgements

We would like to thank David Johnston and the Southampton Biomedical Imaging Unit for their assistance regarding the use of the SEM and CLSM.

Proteomic data is available at the following: https://eprints.soton.ac.uk/435043/

## Notes

**Source(s) of Support** This work was funded by a Biotechnology and Biological Sciences Research Council CASE Studentship award in partnership with Helperby Therapeutics, (BB/L016877/1). Instrumentation in the Centre for Proteomic Research is supported by the BBSRC (BM/M012387/1) and the Wessex Medical Trust.

**Declarations of Interest** YH and ARMC are shareholders in Helperby Therapeutics Group plc. YH is the Director of Research and ARMC is a company founder and the Chief Scientific Officer.

https://eprints.soton.ac.uk/435043/

